# Assessing the hydromechanical control of plant growth

**DOI:** 10.1101/2023.09.21.558781

**Authors:** Valentin Laplaud, Elise Muller, Natalia Demidova, Stéphanie Drevensek, Arezki Boudaoud

## Abstract

Multicellular organisms grow and acquire their shapes through the differential expansion and deformation of their cells. Recent research has addressed the role of cell and tissue mechanical properties in these processes. In plants, it is believed that growth rate is a function of the mechanical stress exerted on the cell wall, the thin polymeric layer surrounding cells, involving an effective viscosity. Nevertheless, recent studies have questioned this view, suggesting that cell wall elasticity sets growth rate or that uptake of water is limiting for plant growth. To assess these issues, we developed a microfluidic device to quantify growth rates, elastic properties, and hydraulic conductivity of individual *Marchantia polymorpha* plants in a controlled environment with a high throughput. We characterized the effect of osmotic treatment and of abscisic acid on growth and hydromechanical properties. Overall, the instantaneous growth rate of individuals is correlated to both bulk elastic modulus and hydraulic conductivity. Our results are consistent with a framework in which growth rate is determined primarily by elasticity of the wall and its remodelling, and secondarily by hydraulic conductivity. Accordingly, the coupling between chemistry of the cell wall and hydromechanics of the cell appears as key to set growth patterns during morphogenesis.

## 1. Introduction

Plant growth and morphogenesis are complex multiscale phenomena where both (bio)chemistry and (bio)mechanics play a fundamental role. While there is little questioning that mechanical forces are involved in growth [1–5], the relevance of different hydromechanical (hydraulic and mechanical) properties in growth is still subject to diverging views, notably the role of water conductivity and elastic deformations of cells and tissues [5–7]. The high concentration inside plant cells creates a differential of water chemical potential that drives water into the cell [6], creating a high mechanical pressure called turgor that is resisted by the stiff cell wall, the polymeric thin layer around cells. The balance between turgor pressure and wall resistance sets growth rate of cells and tissues [1–5,8]. Uptake of water is dependent on the plant water conductivity, a measure of the amount of water that crosses a unit surface of the plant per unit time, driven by a unit drop in water potential (volumetric chemical potential of water) across the surface (and so expressed in m.s^-1^.Pa^-1^). Whether growth may be limited by hydraulic conductivity has been debated given the experimental data collected on specific organs or full plants [6,7]. Recently, there has been a renewed interest in this question as well as in the role of turgor pressure in growth [9]. Experimental and theoretical studies suggested that water conductivity of cells limits their growth and plays a role in the fine tuning of turgor pressure [10,11]. In this study, we further examine the relation between hydraulic conductivity and growth rate of the tissue.

The wall’s ability to deform under pressure controls the growth dynamics along with the regulation of turgor pressure. This ability depends on mechanical properties (elasticity, viscosity…) of the wall, remodelling of its architecture, and changes in its composition. Attempts to model wall expansion fall into different frameworks, most of them considering either stress (force) or strain (deformation) as the main determinant of growth rate [12]. Stress (***σ***) is a measure of the physical forces applied on a material, corresponding to a force per unit surface applied to the sample and expressed in Pascals (Pa). The strain is a measure of said material’s deformation under stress, corresponding to a relative increase of dimensions of the sample. In the case of an elastic material, the elastic modulus is the proportionality constant between stress and strain. In this study we provide experimental evidence that elastic strain and material elasticity are key to understanding growth regulation.

Most frameworks on growth regulation consider the relative growth rate ***G*** (expressed in s^-1^), also known as strain rate or relative expansion rate, which is the relative rate of increase in dimensions. In the present study we use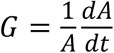, with *A* the plant surface area and *t* the time. In such frameworks, the relative growth rate the cell wall depends on stress in the wall. This is typically the case in the classical Lockhart model [13] where ***G*** is proportional to stress **σ** in excess of a threshold ***Y*** (yield stress), the proportionality factor being the extensibility **Φ** (expressed in Pa^-1^.s^-1^), which may be interpreted as the inverse of an effective viscosity:

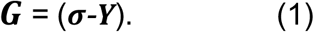

This seminal description spawned a whole class of viscoplastic models of plant growth [14–17]. Some of those models include the elastic modulus in their mechanical parameters but their final formulations make the modulus dispensable to describe steady state growth, ending in description where only viscosity (or extensibility) is relevant. Experimental work was conducted to assess the validity of Lockhart’s model [18–20] as well as to understand further the biological basis of the relevant parameters [21,22] and get an estimation of their value [19,23]. However, it is important to note that the results of the aforementioned work could also be interpreted outside of this stress-derived framework, because stress is proportional to elastic strain, a measure of the reversible deformation of the sample under stress.

Notably fewer theoretical studies considered elastic strain as the relevant mechanical variable to set growth rate [24,25]. Nevertheless, mechanistic considerations on remodelling of the cell wall led to chemo-mechanical models of cell wall expansion that are compatible with such strain-based model of growth [24,26]. In this framework cell wall expansion rate is proportional to elastic strain and so is inversely proportional to elastic modulus (for a given turgor pressure). The proportionality prefactor is a time scale that may be related to the chemistry of the cell wall. Recent experimental work led to progress in the measurement of the cell wall mechanical properties and in particular of its elastic modulus [27]. Some of these studies hinted to the importance of elasticity and elastic strain in the regulation of growth in the shoot apex of flowering plants. Organ outgrowth was found to be dependent on cell wall remodelling (pectin methyl-esterification in this case), which was associated with a reduction in elastic modulus [28,29]. Within the shoot apex, domains growing at a slower rate were found to have smaller elastic strain based on osmotic treatments [30] and higher elastic modulus based on atomic force microscopy [28,31,32]. In the roots of several species, fast growing regions were found to have a lower modulus [33].

The difference in modelling between a viscous approach (based on extensibility) and an elastic approach (based on elastic modulus and a chemistry derived time constant) may seem small as they end up with the same mathematical description of growth dynamics. They however differ greatly in terms of underlying mechanisms. Overall, available experimental results do not strongly discriminate between elastic strain and stress as the main determinant of growth rate. This is in part due to the difficulty of measuring mechanical properties and growth characteristics of the same individuals with a high throughput.

Altogether, it appears that the hydromechanical basis of growth control in plants is far from being fully understood. In order to make progress, we chose *Marchantia polymorpha* as a model organism, using a microfluidic system to grow small plants in the same conditions. While Marchantia plants provide large numbers of genetically identical progeny by vegetative propagation, microfluidic chips enable the observation of a high number of individuals over time [34–36]. We developed a microfluidic-based approach to characterize hydromechanical properties, giving us access to growth rate, elastic modulus, and hydraulic conductivity of the same individual plants with high throughput.

## 2. Results

### A. High-throughput characterization of growth and hydromechanics in controlled conditions

#### Growth

To observe in parallel a large number of individuals in the same conditions, we designed a microfluidic chip with V-shaped traps (Fig. 1-A) to immobilize 50-100 small *Marchantia* plants (gemmae) during 30h. As gemmae are dispersed by rain in the field, they are prone to be bound by a film of water after they have left the mother plant and so are likely tolerant to immersion. In our system, gemmae were immersed in liquid growth medium (see methods for details), which was continuously renewed by slow injection in the chip during the experiments. We first optimized the collection of gemmae from the mother plant and ensured that growth of the gemmae was robust to the choice of experimental conditions, notably flow rate and immersion (Fig. S1). We tracked gemmae growth with a microscope (Fig. 1-B) and retrieved their area using image analysis (see methods). We observed an early phase with no visible growth, followed by a phase of approximately exponential growth (Fig. 1-C).

**Figure 1.**
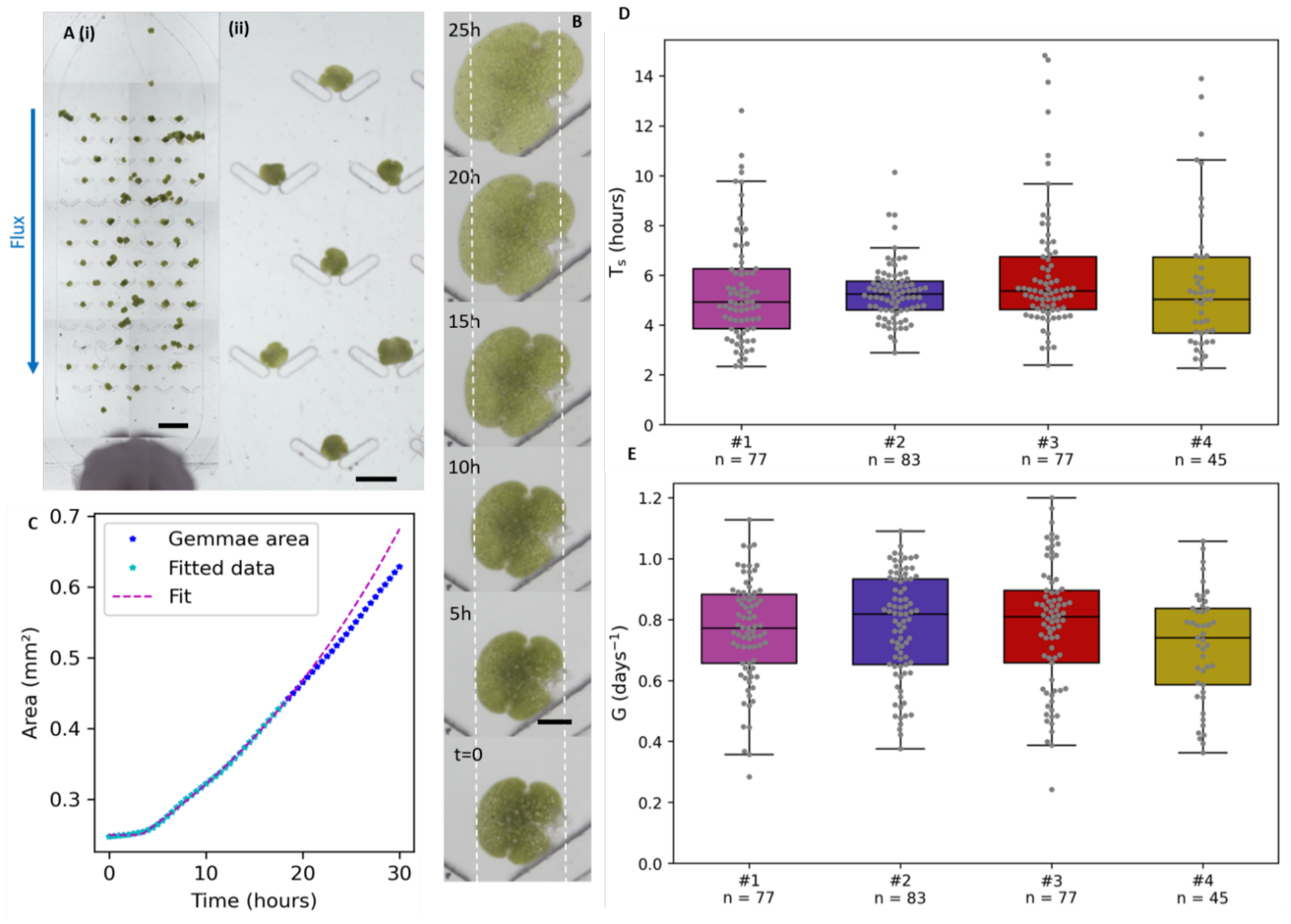
Growth in a microfluidic chip. **A. (i)** Microfluidic chip design with V-shaped traps holding several gemmae (small Marchantia plants) for parallel observation. Scale bar: 5 mm. **(ii)** Zoom on traps. Scale bar: 1mm. **B**. Growth of a gemma in time. The white dashed lines serve as a marker of initial size. Scale bar: 250µm. **C**. Area vs. time curve for the gemma represented in B (blue). The magenta dashed line is a delayed exponential fit. Fitted points (until 15h after growth start) are shown in light blue. **D**. Values of growth start delay for 4 independent experiments. **E**. Values of average growth rate during 15h for the same experiments as in (D).

We characterized this dynamics with two parameters, ***T***_***s***_ and ***G***, obtained by a delayed exponential fit of each growth curve up to 15h of exponential growth (see methods). ***T***_***s***_ is the time between the immersion of plants in solution (which is analogous to the start of imbibition for seeds of seed plants) and the start of exponential growth (analogous to the onset of growth during seed germination). ***G*** is the increase rate of the exponential.

In total we quantified growth for n = 282 gemmae in N = 4 independent experiments (Fig. 1-EF). The median growth start time ***T***_***s***_ over all experiments was 5.20 hours (4.93 to 5.38 hours for individual experiments medians). The variability (expressed as relative average absolute deviation, see Materials and Methods for more details on statistical analysis) was 28% over the complete distribution (from 16% to 40% in individual experiments) showing a highly variable time of growth start. The relative growth rate ***G*** had a median of 0.78 days^-1^ (0.74 to 0.82 days^-1^) with a variability of 19% (17% to 20% in individual experiments) indicating that the growth rate of gemmae was also quite variable. For both measurements the variability in single experiments was higher than the difference across experiment medians (∼10%), underlining the fact that the overall variability is representative of the variation between gemmae and not an artifact of data pooling. Such level of variability within a single sample is reminiscent of seed germination time, which can be largely variable even for seeds placed in identical conditions [37].

#### Bulk elasticity

The microfluidic system that we built also enables rapid (∼1min) change of the medium in which gemmae are developing. We took advantage of this to design a method to measure the mechanical properties of gemmae. This method was based on osmotic steps during which the control growth medium is replaced with a hyperosmotic medium (control medium supplemented with 100mM mannitol, for a total osmolarity of about 130mM). This treatment drove water out of plants and leads to the contraction of the gemmae by a few percent in area that can be observed and measured through image analysis (Fig. 2-AB). The reverse step was applied 15 minutes later by refilling the chip with the control medium (∼30mM of solutes) and the re-inflation of the gemmae was observed and quantified in the control medium as well. We used the curve associated with these changes in area to extract a bulk elasticity modulus from the deflation (hyper-osmotic step) and to quantify deformation irreversibility from the re-inflation in control medium (see Materials and Methods for details). The first osmotic step appeared to stop gemmae growth during the 15 minutes of treatment but for the return step and reinflation we were led to account for growth of the gemmae to get a better measurement of recovery strain (Fig. 2-B and Materials and Methods for details on the fitting). We could thus quantify the elastic modulus (Fig. 2-C) as well as the irreversibility of the deformation between the two steps (Fig. 2-D). In this experiment (n = 63) we obtained a median bulk elastic modulus of 16.6 MPa with 14% of variability. The distribution of deformation irreversibility had a mean of 107.7% (p = 5.0 10^−10^ in comparison to 100% that corresponds to full reversibility). We speculate that this slight irreversibility might be due either to potential damage to the cell walls caused by the initial step and contraction, or to plant reaction during the 15 minutes between the osmotic step and the return to normal medium. Finally, we assessed the robustness of the quantification of modulus and irreversibility by making two consecutive measurements separated by 30 min (Fig. S2-AB). We found relatively small variations of the two quantities, suggesting that the first measurement does not strongly perturb the state of gemmae.

**Figure 2.**
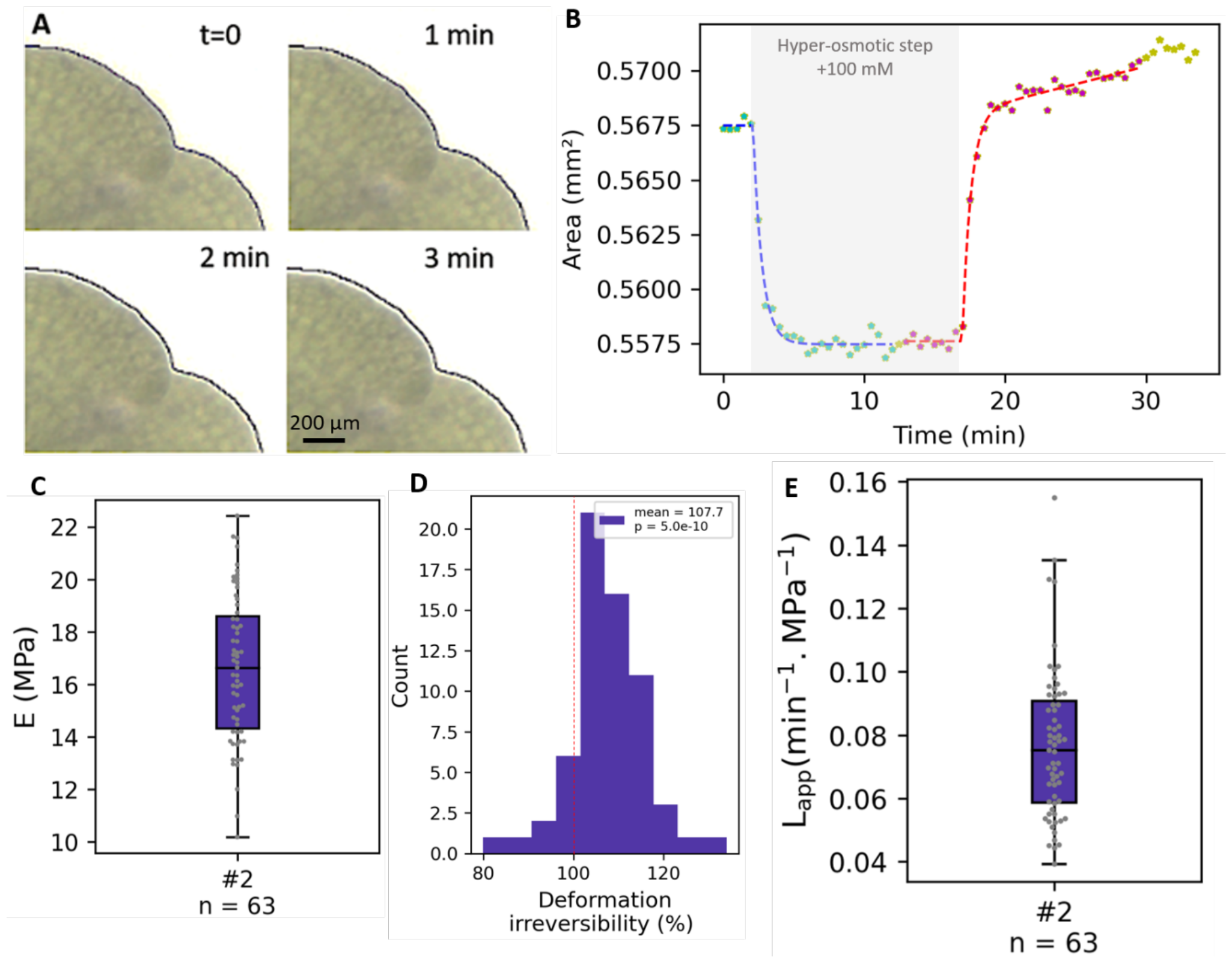
Measuring hydromechanical properties with osmotic steps. **A**. Illustration of the deformation of a gemma subjected to an osmotic step (100mM mannitol). A portion of a gemma is shown; the black line represents the contour of the gemma at t=0 to underline the deformation. At t = 2 and 3 minutes the thin white line between the black line and the gemma corresponds to the contracted area. **B**. Dynamics of the area of a gemma when subjected to an osmotic step followed by a return to control medium. The light blue dots are fitted by the blue dashed line to obtain the elastic modulus and conductivity, and the mauve dots are fitted with the red line to obtain elastic reversibility. **C**. Distributions of bulk moduli. **D**. Distribution of irreversibility levels. The red dashed line illustrates the complete reversibility to which the distribution mean is compared. **E**. Distribution of apparent conductivity.

To get a better quantification of gemmae elasticity and its variability we pooled this experiment (Fig. 2-CD) together with the controls of the different treatments discussed hereafter (Fig. 4), yielding N = 6 experiments with n = 222 individual plants. The median bulk elastic modulus was 13.95 MPa (11.9 to 16.6 MPa in individual experiments) and the variability was 16% (11% to 14% in individual experiments). Contrary to what we observed for ***T***_***s***_ and ***G***, here the global variability came from the difference between experiments more than from internal variability. Regarding deformation irreversibility, individual experiments ranged from 101.7 to 107.7% of irreversibility, and were statistically different from 100% in 3/6 experiments, consistent with a limited impact of osmotic steps on gemmae. The standard error of those distributions was between 1.0 and 2.8%.

Overall, these measurements are coherent and show that our method is robust. The observed low level of deformation irreversibility also supports the definition of a bulk elastic modulus for gemmae.

#### Hydraulic conductivity

Using the dynamics of deflation (Fig2-B), we also designed a method to extract the apparent hydraulic conductivity, ***L***_***app***_, of the plant, based on modified Lockhart equations (see Materials and Methods). Note that this is an individual-level conductivity accounting for cell membranes, plasmodesmata, cell wall, and cuticle. It is proportional to the classical hydraulic conductivity, ***L***, involving gemma thickness, ***H***: ***L***_***app***_ *=* **2*L*/*H*** (see Material and Methods) and has units of inverse time and pressure (e.g. min^-1^.MPa^-1^). We provided here values of apparent conductivity because we could not measure the thickness of individual gemmae in the chip. In the first experiment (Fig2-E, n=63) we obtained a median apparent conductivity ***L***_***app***_ of 0.08 min^-1^.MPa^-1^ with 24% of variability (n=63). When pooling all controls we got a median of 0.09 min^-1^.MPa^-1^ (0.08 to 0.11 in individual experiment) with 22% of variability (10% to 24% in individual experiments). This median corresponds to a water conductivity, ***L***, of the order of 7.10^−8^ m.s^-1^.MPa^-1^, using the typical thickness of the gemmae (***H*** ∼ 100µm from hand-made sections). Like for growth rate, the variability of individual experiments was higher than the variability between experiments median (∼15%), here again indicating a high internal variability of the apparent hydraulic conductivity. We also measured hydraulic conductivity twice over 30 min (experiment mentioned above, Fig. S2-C) and found relatively small variations of conductivity, suggesting that the first osmotic step does not significantly perturb the plants.

### B. Characterisation in perturbed conditions

#### Growth

To further probe the relation between modulus and growth rate, we used our system and analysis pipeline to quantify variations due to classical treatments. We considered changes of growth dynamics caused by external physico-chemical constraints or perturbed signalling. For each condition we used two microfluidic chambers in parallel, one control and one perturbed condition, with a common gemmae pool split between the chambers, and repeated the experiment N = 3 times. The aggregated results are presented in Fig. 3 and the detailed results per experiment are available in Fig. S3.

**Figure 3.**
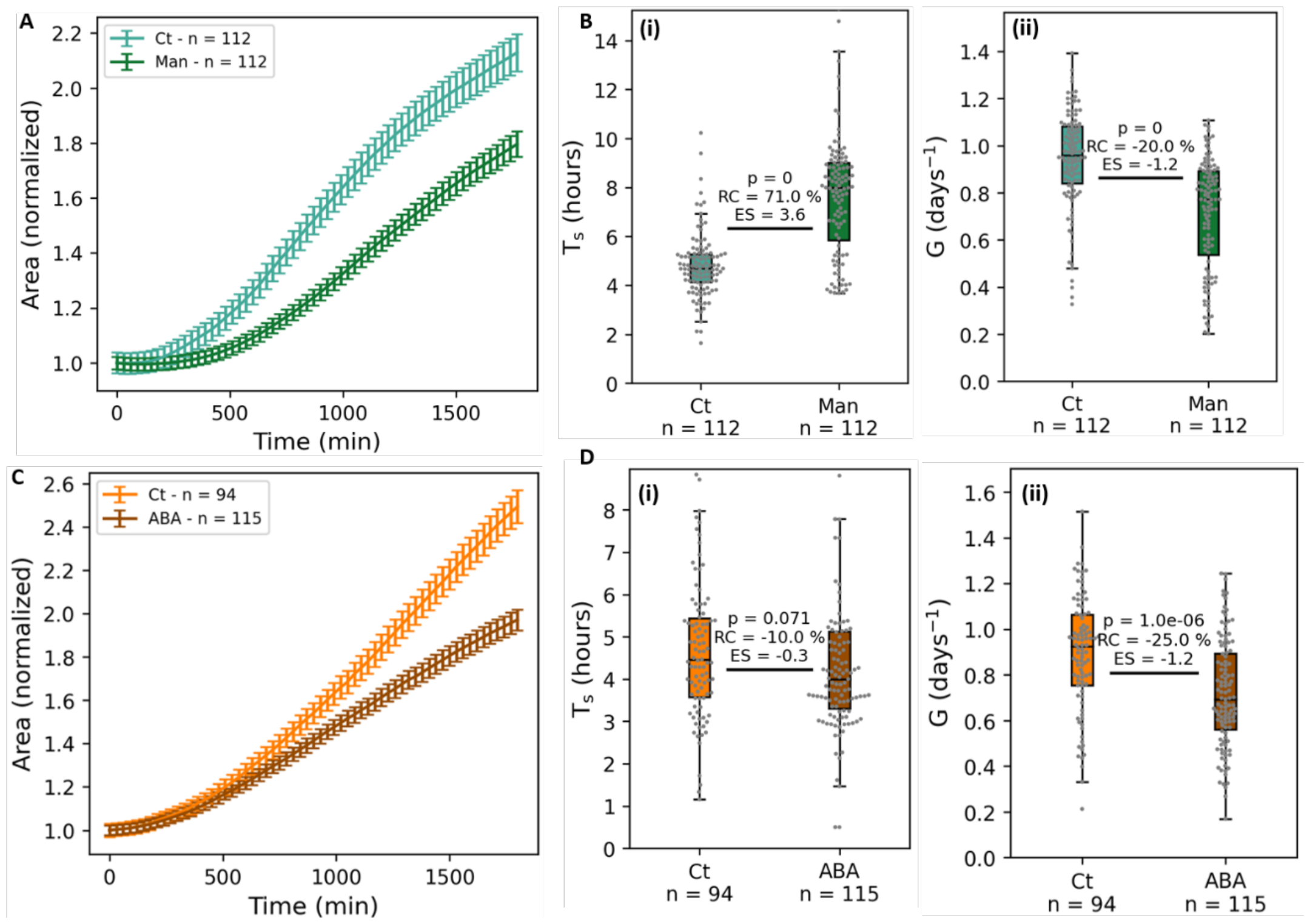
Growth in perturbed conditions. **A**. Area vs. time curves illustrating growth in hyperosmotic medium — reference medium supplemented with 100mM mannitol (shortened Man) — compared to reference medium (control, Ct). Each curve represents average and standard error for all individuals pooled from N = 3 experiments. The area curves are normalized by the initial (t=0) value for better comparison between conditions, see Fig. S3 for detail by experiment **B**. Quantification of growth start time (i) and growth rate (ii) for N=3 experiments in hyperosmotic medium. **C**. Area vs. time curves illustrating growth with abscisic acid (ABA, 3.5µM) treatment compared to reference medium (control, Ct). Each curve represents average and standard error for all individuals pooled from N = 3 experiments. The area curves are normalized by the initial (t=0) value for better comparison between conditions, see Fig. S3 for detail by experiment **D**. Quantification of growth start time (i) and growth rate (ii) for N=3 experiments with ABA.

We started with hyperosmotic treatment because this classically reduces growth rates. Gemmae were observed in control medium permanently supplemented with 100mM of mannitol (Fig. 3-A). This treatment significantly reduced the growth rate of gemmae, with a relative change RC = - 20% when compared to control (p < 10^−10^) and an effect size (see methods) ES = -1.2, as expected. Another effect of growing in a hyper osmotic environment (Fig. 3-AB) was a largely delayed growth start with a relative change RC = -71% (p < 10^−10^) and an effect size ES = -3.6 indicating a strong change.

We then performed chemical treatment using a medium complemented with the abscisic acid (ABA) phytohormone. ABA is produced when water is lacking and enables plants to adapt to water stress [38,39]. The response to ABA was proposed to induce a delay in dormancy exit of Marchantia gemmae assessed from rhizoids emergence [40]. Based on these previous studies, we used a concentration of 3.5µM ABA. Unexpectedly, this treatment (Fig. 3-CD) had a small, non-significant decreasing effect on the growth start time (RC = -10%, ES = -0.3, p = 0.071). However, it reduced growth rate like the hyper-osmotic treatment (RC = -25%, ES = -1.2, p = 10^−6^), consistent with ABA-treated plants being smaller in [39].

Overall, those perturbations lead to changes that are easily quantifiable with our system, in particular thanks to the observation of large numbers of individuals in identical conditions.

#### Bulk elasticity

We then moved on to quantify the effect of those treatments on the hydromechanical properties of the gemmae (Fig. 4). We performed osmotic steps on gemmae that grew for 30h either in +100mM hyperosmotic medium or with 3.5µM ABA.

For the gemmae that grew in hyperosmotic medium (Fig. 4-A & S4-AB, N = 3, n = 95 & 80 for control and treated sample, respectively) we observed a decrease of the bulk elastic modulus, *E*, when compared to their control (RC = -29%, ES = -2.0, p < 10^−10^). In addition, we saw a small decrease of deformation irreversibility in the treated sample (RC = -4%, ES = -0.7, p = 2.0 10^−4^) (Fig. S4-A).

Regarding the effect of ABA treatment (Fig. 4-B & S4-DE, N = 2, n = 64 & 78) we observed a small, non-significant, decrease of elasticity (RC = -6%, ES = -0.2, p = 0.067). The irreversibility was unaffected by ABA treatment (Fig. S4-D).

#### Hydraulic conductivity

Finally, we examined the apparent hydraulic conductivity ***L***_*app*_. Hyperosmotic medium (Fig. 4-C & S4-C, N = 3, n = 95 & 80) slightly increased conductivity (RC = +12%, ES = 0.6, p = 0.014) in comparison to control. For ABA treated gemmae (Fig. 4-D & S4-F) we observed a very large increase of ***L***_***app***_ (RC = +60%, ES = 3.4, p = 5.410^−10^) when compared to control.

### C. Relation between growth rate and hydromechanical properties

We then asked if there was a relationship between the growth rate of a gemma and its hydromechanical properties. The system that we designed is well suited to answer such a question, as osmotic steps can be performed on a gemmae population after observation of its growth. It is thus possible to obtain a bulk elastic modulus and conductivity for gemmae whose growth dynamic has previously been characterized. When first looking at the relation between growth rate ***G*** and hydromechanical properties (***E***, and ***L***_***app***_) in controlled conditions, we found low correlations, significant for modulus ***E*** (CC = -0.197, p = 0.008) and non-significant for conductivity ***L***_***app***_ (CC = 0.135, p = 0.071) (Fig. S5-B(i)C(i)). As elasticity is measured from a fast deformation (a few minutes) compared the area doubling time (several hours), we sought a more instantaneous characterization of growth, as close as possible in time to the measurement of elasticity. For this we measured the instantaneous growth rate 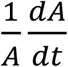 from the growth curve and took its average value over the last 4 time points (∼2 hours, see Fig. S5-A) to define ***G***_***inst***_ as the growth rate just before the step. We found a correlation coefficient of -0.503 (Spearman’s rho, p < 10^−10^) (Fig. 5-A) between ***G***_***inst***_ and the modulus ***E*** and a correlation coefficient of 0.318 (Spearman’s rho, p = 2.10^−5^) between ***G***_***inst***_ and the conductivity ***L***_***app***_ for all controls pooled.

**Figure 4.**
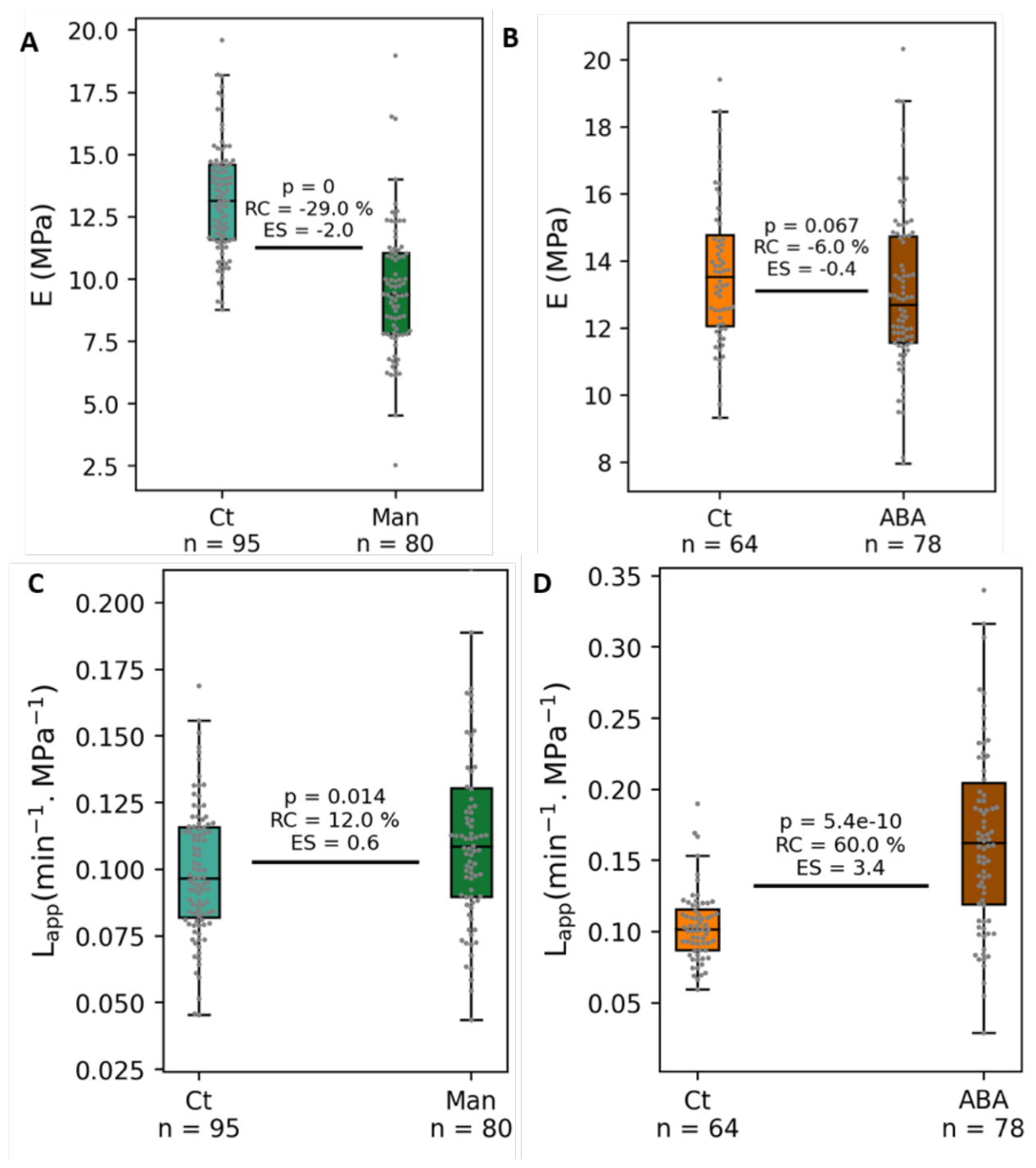
Hydromechanical properties in perturbed conditions. **A**. Elastic moduli for control (Ct) and mannitol (Man) treated gemmae. **B**. Elastic moduli for control (Ct) and abscisic acid (ABA) treated gemmae **C**. Apparent conductivity for control (Ct) and mannitol (Man) treated gemmae. **D**. Apparent conductivity for control (Ct) and abscisic acid (ABA) treated gemmae.

**Figure 5.**
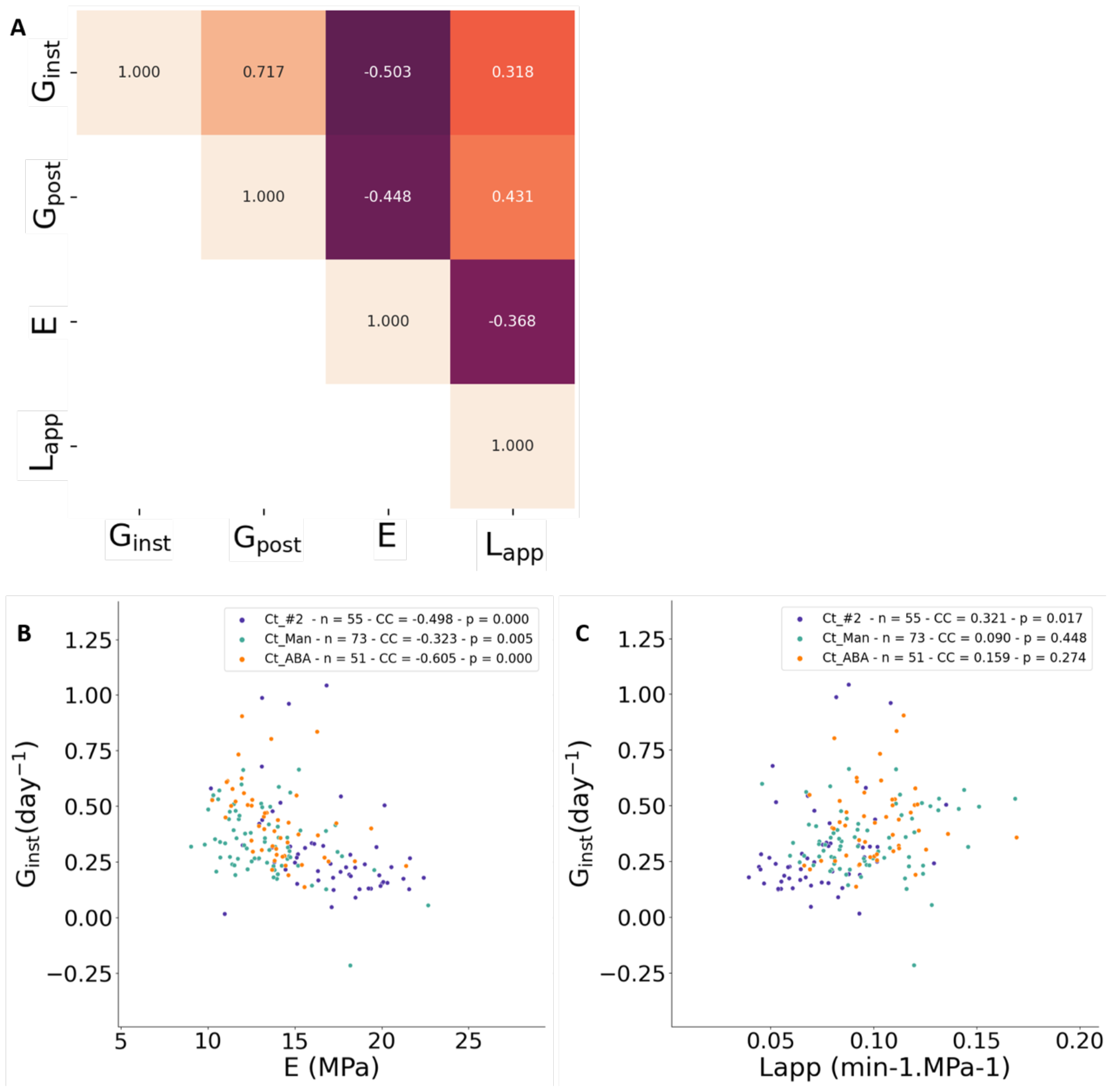
Link between hydromechanical properties and growth: bulk modulus E, apparent conductivity L_app_, instantaneous growth rate G_inst_, and growth rate avec the steps G_post_. **A**. Correlation table for all control experiments pooled. All correlations have p < 10^−3^. **B**. Relation between G_inst_ and E for all the different controls. **C**. Relation between G_inst_ and L_app_ for all the different controls.

We note that these correlations are based on gemmae for which both growth and elasticity/conductivity are properly quantified, reducing the number of gemmae to n = 179, lower than those observed growing (n = 289, Fig. 1-DE & Fig. 3) and those characterized mechanically (n=222, Fig. 2-C & Fig. 4). This is due to the fact that performing osmotic steps implies a fast flow of medium that can push a few gemmae away from the observation field.

In the case of elasticity, we found a clear decreasing relationship between elastic modulus ***E*** and instantaneous growth rates ***G***_***inst***_ (Fig. 5-B). This significant correlation (p < 10^−10^) showed that being softer goes together with faster growth, without indicating any causality of effect between those two parameters. The fact that the correlation is improved when considering instantaneous growth rate suggested that the mechanical properties and instantaneous growth rate change together in time during gemmae development. In the case of conductivity, we found a positive, significant relationship (p = 2.10^−5^) between ***L***_***app***_ and ***G***_***inst***_ (Fig. 5-C). Given the definition of ***L***_***app***_ we wondered about the influence of gemmae thickness ***L*** on this correlation. Noting that growth rate correlates with area at 30h and that area correlates with gemmae thickness, the positive correlation between ***L***_***app***_ *=* ***L***/***H*** and growth rate implies that the conductivity ***L*** correlates positively with ***G***_***inst***_. Accordingly, the hydraulic conductivity is higher in plants that grow faster.

We then further examined potential causal relations between these variables, keeping in mind that causality cannot be demonstrated only using correlations. To do so, we considered the relation between hydromechanical properties (***E*** and ***L***_***app***_) and the growth rate after the osmotic step (***G***_***post***_, computed from the end of the re-inflation curve, see methods). We observed a slightly weaker correlation between ***E*** and ***G***_***post***_ than with ***G***_***inst***_ and a stronger correlation between ***L***_***app***_ and ***G***_***post***_ than with ***G***_***inst***_ (Fig. 5-A). For each parameter (***E*** and ***L***_***app***_) we compared correlations with ***G***_***post***_ and with ***G***_***inst***_ to see if the difference between them was significant (see Materials and Methods). We found that the differences are not significantly different in the case of modulus ***E***. However, the correlation between ***L***_***app***_ and ***G***_***post***_ is significantly stronger than the one with ***G***_***inst***_. Given that we first measured ***G***_***inst***_, then ***L***_***app***_, then ***G***_***post***_, this suggested that the conductivity may predict the growth rate.

## 3. Discussion

We developed a microfluidic system to study the early growth of *Marchantia* gemmae in controlled environments and with high throughput. The environmental control given by microfluidics allowed us to perform osmotic steps on growing gemmae and thus directly link their hydromechanical properties to their growth rate. In the course of 12 independent experiments, we quantified the growth of 715 individuals and the hydromechanical properties of 426 individuals. The values that we found for the bulk elastic moduli are in the lower range of moduli measured for plant tissues [1] but have the same order of magnitude as those measured in several bryophytes [41]. The observed small irreversibility, with a higher deformation after the return to control medium could also be a sign of stored growth. This phenomenon qualifies the continuation of cell wall synthesis without visible change in dimensions during phases of low turgor due to osmotic treatment, followed, upon restauration of turgor, by an enhanced growth rate that yields the same plant dimensions as if growth had not stopped by the treatment [20,42,43]. Bespoke experiments would be needed to test this idea. Finally, the values of water conductivity (∼7.10^−8^ m.s^-1^.MPa^-1^) are in the same range as the conductivity measured in roots of flowering plants [44–46].

For 179 individuals among the control experiments we had access to both hydromechanical properties and growth rate, which allowed us to show that there is a correlation between bulk elasticity of the gemmae and their instantaneous growth rate, in line with previous reports of stiffer regions of the shoot apical meristem of tomato growing slower [30] and softer regions of roots growing faster [33]. A similar correlation was recently observed at cell wall level in Marchantia [47]. The weaker correlation between hydraulic conductivity and growth rate suggests that growth rate is partially limited by the plant water conductivity. However, this correlation arises when considering a very large number of individuals and is not necessarily significant in individual experiments where the median conductivity is larger (Fig. 5-C). This suggests that water conductivity of Marchantia in control conditions is in a critical range where it does not always limit growth. This critical range could be relevant in terms of growth control as it only takes a small change of conductivity to either slow down growth or to release the limitation by hydraulic conductivity. This critical range may also explain diverging results and the ensuing debate about the role of water conductivity in plant growth [6,7]. In addition, given the observed variability of growth rate in time (see Fig. S5-A for a representative curve), and the variability of hydromechanical properties across experiments, our results suggest that the regulation of instantaneous growth rate is directly linked to hydromechanical properties of the plant.

When subjecting gemmae to water-stress by the means of a permanent hyper-osmotic environment, a delay in growth start as well as a reduced growth rate were observed, consistent with classical observations on plant growth in water-stressed conditions [48,49]. We also found gemmae to be softer after having grown in hyperosmotic medium. This was observed for the unicellular algae *Chlorella emersonii* [50] and is in line with early reports of osmotic treatment-induced extensibility variation in maize [42,51], where the increase of extensibility might be interpreted as a decrease of elastic modulus. Our results significantly strengthen those observations thanks to a tight control of experimental conditions and a large number of individuals probed. Water conductivity is slightly increased by this treatment. Even though osmotic stress generally decreases root hydraulic conductivity [44], is has been shown to increase it in some accessions [46]. Taken together the increase of conductivity and decrease of elastic moduli suggest that the reaction of the plant to osmotic stress is to adjust its hydromechanical properties to increase growth rate (Fig. 5), partially compensating the growth rate reduction caused by the reduced osmotic differential.

When gemmae were treated with abscisic acid (ABA), we observed a reduction in growth rate, in line with previous observation of ABA treatment on plant growth [52]. Surprisingly ABA treatment had no visible effect on the timing of growth start. Because ABA treatment has previously been reported to delay dormancy exit in *Marchantia polymorpha*, as measured by the first appearance of rhizoids [40], this result raises the question of the appropriate criterion for dormancy exit definition. We measured a growth start around 5h while the time of rhizoids appearance is rather at the timescale of a day. We may postulate that the delay in rhizoid appearance would be due to a slower growth that would make rhizoids reach the gemmae periphery later, rather than with a delayed dormancy exit. Regarding hydromechanical properties, growth in ABA medium did not affect the bulk elasticity of gemmae, but greatly increased their water conductivity. Similar results have been found in roots in response to ABA [45]. This shows that ABA plays a role in water stress regulation [51] by modulating water conductivity rather than mechanical properties. If we consider the previous observation that ABA reduces cell wall extensibility [52], this suggests that ABA also affects the dynamics of wall remodelling.

Taken together with previous work [24], our results on growth and mechanics support a modelling framework where, in stationary conditions, the relative growth rate depends on ***E*** as

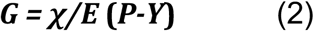

where *χ****/E*** is the analogue of the extensibility ***Φ*** (in Pa^-1^.s^-1^) in Lockhart’s model (Eq. 1 in the introduction), ***E*** is the elastic modulus (in Pa), ***P*** is the turgor pressure (in Pa) and *χ* (in s^-1^) is a chemical rate corresponding to cell wall remodelling, and ***Y*** is a pressure threshold (in Pa) for expansion to occur. It can also be expressed as

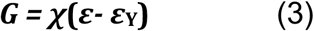

with ***ε = P/E*** the elastic strain and ***ε***_**Y**_= ***Y/E*** the deformation threshold for expansion to occur. Equations (1) and (2) describe the same phenomenon, but with a different mechanistic interpretation. When Ortega derived his augmented growth equation from a Maxwell linear viscoelastic model, he considered the extensibility to be the inverse of the wall viscosity [54], but recent studies show that the cell wall mechanics has a plastic component [55], and found fast relaxation times associated with overall wall deformation [28] or local deformation within the wall [56] that are orders of magnitudes away from the plant growth rate. Overall, a mechanistic interpretation of such viscosity is lacking. We propose to interpret extensibility ***Φ*** = *χ****/E*** as the capacity of the wall to remodel itself at a rate ***χ*** when under an elastic deformation ***ε*** defined by ***P*** and ***E***, as the expression in (3) suggests. This leads to a reinterpretation of plant growth modulation by osmotic and ABA treatment: osmotic treatment reduces the modulus ***E*** [42] and thus increases extensibility ***Φ***, while ABA treatment reduces remodelling rate so ***Φ*** decreases as well [52], assuming other parameters constant. This framework is also in better agreement with studies associating elastic strain and growth rate [30].

We may include equation (2) into the full Lockhart-Ortega equations describing the relative area increase in time of a gemma:

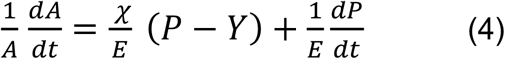

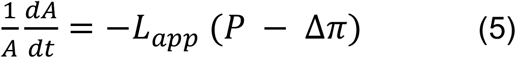

with *A* the area of the gemma and *Δπ* the osmotic pressure differential between the inside and outside of the gemma. Equation (4) represents the contributions of growth and elastic deformations to changes in area, while Equation (5) represents the changes in area associated with water uptake. When osmotic pressure is constant 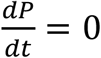, (4) and (5) can be combined by elimination of P, yielding

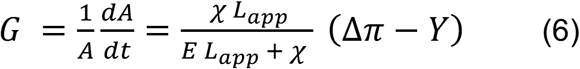

Equation (6) recapitulates our results (see Fig. S6 for a schematic drawing): when conductivity,

***L***_*app*_, is very high growth rate is proportional to *χ****/E*** and conductivity is not limiting for growth. When ***L***_*app*_ is comparable to *χ****/E***, growth rate is negatively correlated with ***E*** and positively correlated with ***L***_*app*_, as observed in our experiments.

In conclusion, we developed an experimental system to investigate growth and hydromechanics in small plants with the ability to control the growth environment. We have shown that both elastic modulus and water conductivity are relevant to growth control and that altered growth conditions cause a change of these parameters. Altogether, we believe that our experimental approach and results will help overcome the challenges raised by modelling growth of cells and organisms [57].

## 4. Materials and Methods

### A. Plant material

The Tak-1 wild-type accession of *M. polymorpha* was used in this study. Plants were grown on medium (0.2%, Gamborg’s B5 media w/ vitamins, Duchefa) mixed with agar (1.2% plant agar, Duchefa). Plant growth took place in a phytotron (Aralab) with 24h lighting (4600 - 5100 lux) at 22°C. The gemmae used for the experiments were taken from branch order 2 gemmae cups (see Fig. S1-A (iv)) of 3-5 weeks old plants.

### B. Chip fabrication

Microfluidic chips were made of PDMS (Sylgard 184) and cast in a brass mould. The mould was fabricated by micro-drilling (50µm drill) to obtain a non-uniform height for the chamber. The PDMS mix (1:10 ratio) was poured in the mould, degassed for 15 minutes in a vacuum chamber and cured at 70°C for 90 minutes. Once cut out from the mould the chip’s inlet and outlet were created using a biopsy puncher. Chip thickness varied from 1mm (inlet and outlet) to 300µm (trap section) and the total chip volume was ∼500µL. The gradient in thickness allowed the gemmae to enter the chip in any orientation without clogging and to be gently guided to a horizontal position in the trap section for observation.

### C. Experiment preparation

The PDMS chip was bonded to a 76x52x1 mm glass slide using a plasma oven (Harrick Plasma, PDC-002). The inlets were connected to glass syringes (10 & 25mL, SGE Trajan) using PTFE tubing of 1.06 mm inner diameter (Adtech) to allow for the passage of gemmae without clogging. The chip was then filled with medium using syringe pumps (Nemesys low pressure module, CETONI). Gemmae were collected in 1.5 mL tubes containing medium with biocompatible surfactant (1/1000 dilution, TWEEN 20, Sigma) to avoid aggregation and sticking to the collection tube and tubing during preparation. They were then transferred to the chip manually using a 5 mL syringe (SGE Trajan). The initial medium was washed away after a constant perfusion of medium is put in place with a flow rate of 8.33µL/min (500µL/hour) and was maintained throughout the experiment. For the osmotic step, the flow rate used was 500 µL/min to ensure a fast replacement of the medium in the whole chip (∼1 min). This flow rate was maintained for the 15 minutes of the experiment to avoid gemmae movement due to a sudden drop in pressure. For the return step and reinflation the same 500 µL/min flow rate and 15 min duration were used with control medium; the reference flow rate was then restored. For the successive osmotic steps (Fig. S2), the same procedure was repeated twice, with a delay of 30 min between the end of the first return to normal medium and the beginning of the second shift to hyperosmotic medium.

For the no flux experiment corresponding to Fig. S1C the initial medium was washed away by a flux of 50µL/min flux for 10 min, before stopping the flux for the remainder of the experiment. For the 48-well plate experiment corresponding to Fig. S1C there was 1mL of medium in each well, with one gemma per well (except for two wells with two gemmae). The plate was kept on the microscope in the same conditions as a chip experiment (same lighting), and without a lid to ensure an ‘infinite’ supply of atmospheric gases for the plants. Manual adjustment of focus was needed because of changes in liquid level in the wells due to evaporation, so that the number of images taken was much smaller than in microfluidic experiments. We found growth of the gemmae to be reduced with no flux, whereas growth in the 48-plate was equivalent to that in microfluidic chips with a reference flux of medium. We concluded that gas and nutrient supply are not limiting growth in control conditions.

### D. Media and treatments

Control medium was 0.2% Gamborg’s B5 media w/ vitamins (Duchefa) in water with pH adjusted to 5.8. Hyper-osmotic medium was composed of control medium supplemented with 100mM diluted solid D-Mannitol (Sigma). The ABA medium was composed of control medium with 3.5µM Abscisic acid. The ABA powder (Merck) was first diluted in DMSO (Sigma) at 30mM concentration before being used to make the ABA medium.

### E. Image acquisition

Images were obtained at a 16X magnification using a Zeiss Axiozoom V16 and the Zen 3.3 software. Images were taken every 30 minutes for growth experiments, and every 30 seconds during osmotic steps. For growth experiments Z-stacks of 3 images, separated by 150 µm, were acquired at each position. The Z-stacks were reduced to a single image using the Zen post processing function ‘Extended depth of focus’ and the ‘variance’ method with default parameters. For osmotic steps experiments there was no Z-stack acquisition to have a faster acquisition rate.

### F. Image analysis

Videos (tiff image stacks) of individual gemmae were cropped from the original field of view using ImageJ. The rest of the analysis was performed using Python (https://github.com/VLaplaud-Biophysics/PaperDataLAPLAUD2023). The gemmae were detected in each image in several steps: First, the white balance of each video was adjusted. Second, a 3-channel thresholding was applied on the HSV (hue, saturation, value) representation of the original RGB image. The threshold values were adjusted manually for each experiment, but were the same for all gemmae analysed within each experiment. Third, morphological operations (binary closing with 5 µm circular elements, followed by binary opening with a 30 µm circular element) were used on the created binary mask to recover the shape of the gemmae. Finally, areas of gemmae were measured on each image from this reconstituted binary mask.

### G. Data fitting

From the image analysis we obtained experimental curves of the gemmae area in time for both the growth and osmotic step experiments (Fig. 1-C and 2-B). For growth curve parameter extraction, we fitted the experimental data with a delayed exponential of the form:

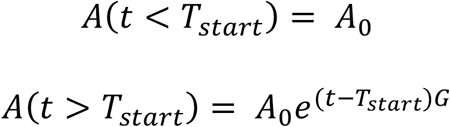

This form takes into account the delayed start and the exponential nature of growth. As we wanted to compare growth rate for a similar time period for each gemma, we used an iterative approach. First, the whole area curve was fitted to obtain a first value of ***T***_***s***_. Then the fit was repeated on a time window stopping 15h after the ***T***_***s***_ previously computed. This yielded a new value for ***T***_***s***_ that defines a new time window ending 15h after it. This process was iterated until the value obtained for ***T***_***s***_ converged with a precision of 0.1 % or reached a maximum 10 iteration in which case the fit was not validated. This ensured a robust definition of ***T***_***s***_ and a value of ***G*** representative of the initial 15h of growth for all gemmae.

For the osmotic steps, each part of the curve (deflation and inflation) was fitted with an evolution function of the form:

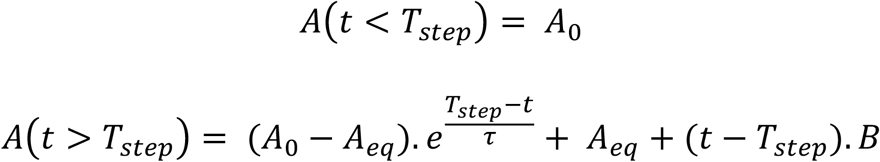

This represents an exponential transition between the two equilibrium states with the addition of a linear component (*B*, in m^2^.s^-1^) to take into account the growth during the step, simply modelled as a linear evolution on a short time scale. The *t – T*_*step*_) factor in both the exponential and linear part of the fit accounted for the fact that it was not experimentally possible to synchronize the imaging of all gemmae in the chip with the exact moment where the hyperosmotic medium reached them. Thus, the beginning of the deflation could happen between two time points, and leaving this time *T*_*step*_ as a fit parameters led to more robust estimates of *τ*.

The elastic modulus was computed from the hyperosmotic step fit using the formula:

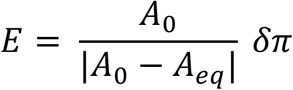

with *δπ* the change in external osmotic pressure.

The deformation irreversibility was computed as

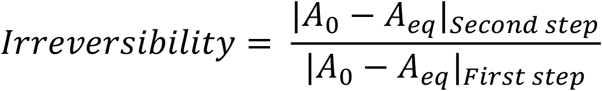

The hydraulic conductivity accounts for cell membranes, plasmodesmata, cell wall, and cuticle. It was extracted from the fit of the gemmae deflation after the hyperosmotic step. We computed it as:

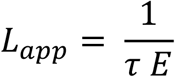

This formula comes from the resolution of a modified Lockhart model adapted to our system:

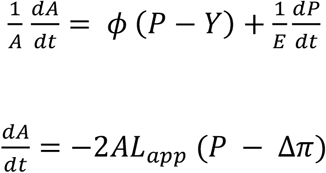

where *A* is the area of the gemma visible on images. The factor of 2 accounts for the upper and lower surface of the gemma. The gemma volume is *V*=*AH*, with *H* gemma thickness, whose variations are neglected considering that *H* is small relative to the other dimensions, and that the number of cell layers is constant. ***L*** *=* ***L***_*app*_*H*/2 is the hydraulic conductivity, *P* the turgor pressure, Δ*π* the osmotic pressure differential between inside and outside the gemmae, *Y* the stress threshold for growth, and E the bulk elastic modulus previously computed. We also assumed (*p* − *y*) *=* 0 as the growth stops between the two steps. We obtained the evolution of the turgor pressure as:

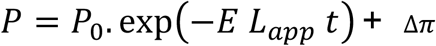

Close to its final value, the area follows the same trend in time as the pressure, allowing us the extract 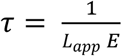 from the fit. Here again, the factor *t* − *T*_*step*_) in the fit ensure that we put a low weight in the fit on the first moments of deflation and thus get a correct value for *τ*.

The growth rate after the step was computed from the re-inflation step as:

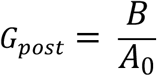

For the ‘instantaneous’ growth rate during growth we computed

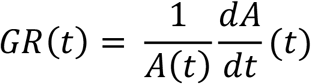

from the experimental data using a first order discretization:

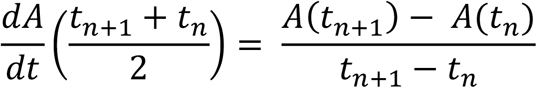

and

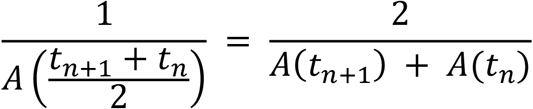

where the *t*_*n*_ are the experimental time points.

### H. Statistical analysis

The variability for data in figures 1 and 2 as stated in the main text were computed as relative average absolute deviation, meaning the average absolute deviation (AAD) of the sample divided by its median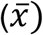:

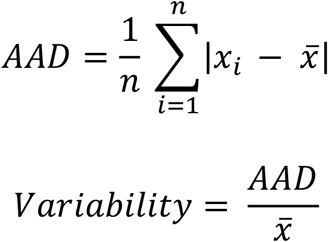

To assess the significance (p-value) of the irreversibility level when compared to 100% we used a one-sided t-test. For statistical analysis and significance (p-values) when comparing two samples the statistics used were non-parametric given the non-normal nature of data distributions in general. When ‘p=0’ is displayed it means that p < 10^−10^. The paired sample comparison in figures S1, S2, 3, S3, 4, and S4 were done using the Wilcoxon ranksum test. For correlations in figures S1, 2, and 4 we used the Spearman correlation coefficient as we could not assume linear relationships between the compared quantities *a priori*.

In addition to p-values we chose to display the Relative Change (RC) and the Effect Size (ES) for each sample comparison. The RC is simply the difference between samples median divided by the control median:

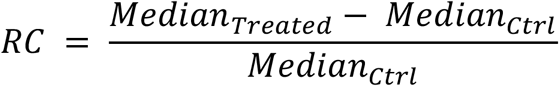

We computed a non-parametric ES as the difference between the two medians divided by the average absolute deviation of the control condition:

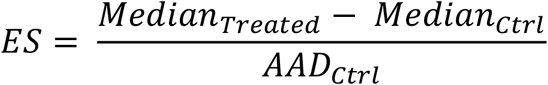

For example, an ES of 0.7 means that the median value of the treated sample is closer to the control median than the average point in the control sample. This gives a way to quantify the difference between two samples while taking into account the variability of the measurements. Computationally we can also understand the ES as the RC between the samples normalized by the variability of the control as defined above:

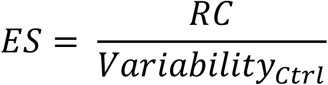

To compare the correlations between ***E*** or ***L***_*app*_ and the growth rates ***G***_***inst***_ or ***G***_***post***_, we used http://comparingcorrelations.org/, an R-based webpage including several methods to compare correlations. In every case stated in section IV all the methods gave the same results.

## Acknowledgments

This work was funded by Agence Nationale de la Recherche (HydroField, grant # ANR-20-CE13-0022-03) and by Ecole polytechnique. The microfluidic device was developed at the microfabrication facility of LadHyx. The authors are grateful to Caroline Frot for her help as the microfluidic engineer of LadHyX.

## Author Contributions

Conceptualization: VL, EM, SD, AB

Methodology: VL, EM, SD

Software: VL

Validation: VL

Formal Analysis: VL, EM, AB

Investigation: VL, ND, EM, SD

Data Curation: VL

Writing - Original Draft Preparation: VL, AB

Writing - Review & Editing: VL, ND, EM, SD, AB

Visualization: VL

Supervision: AB

Project Administration: AB

Funding Acquisition: AB

---

**Figure S1.**
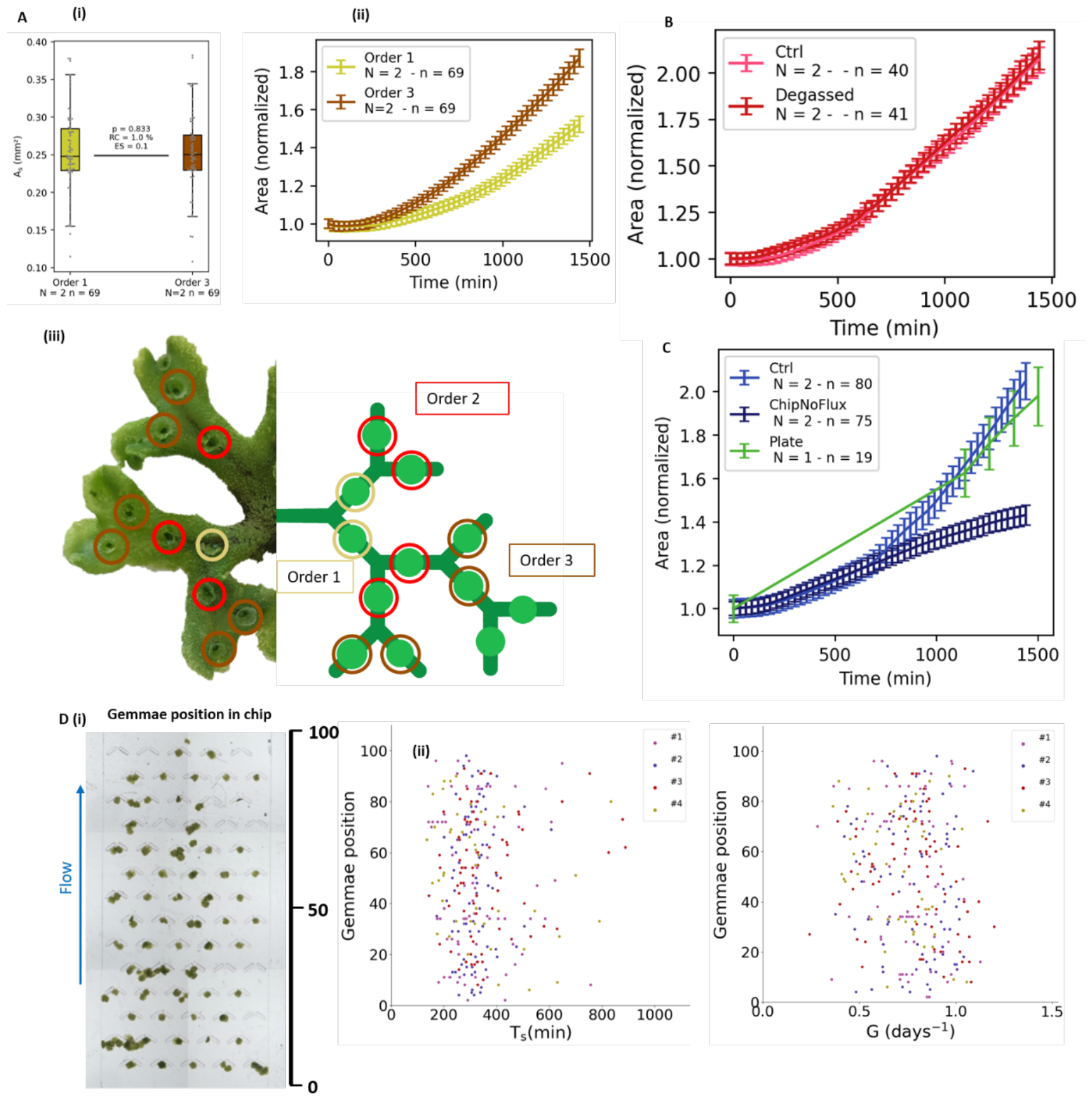
Growth in controlled conditions. **A**. Comparison of growth dynamics for gemmae coming from old (order 1) or young (order 3) cups on the mother plant. (i) Areas at t=0 are similar. (ii) Gemmae from younger cups grow faster than gemmae from older cups. (iii) Illustration of the different branch orders of cups. Light yellow are order 1 (oldest) and yellow are order 3 (youngest) cups. For all experiments other than the one in this figure (S1) gemmae are from order 2 (red) cups. **B**. Comparison of growth dynamics between normal medium and degassed medium. **C**. Comparison of growth dynamics between different flow rates and environments. Ctrl is a chip with reference flow rate, 500µl/h. ‘ChipNoFlux’ is a chip experiment with the flux stopped after 10 minutes to limit gas and nutriment supply. ‘Plate’ is a 48-well plate experiment, with typically one gemma floating in each well initially filled with 1ml of medium, so that neither gas nor nutrient supplies are limiting for growth. **D**. Illustration of gemmae position in the chip (i) and correlation of 36 position with *T*_*s*_ (ii) and *G* (iii) for the data in Fig. 1-DE. The position of the gemmae in the chip does not affect measured growth parameters (Spearman’s rho < 0.15 for both).

**Figure S2.**
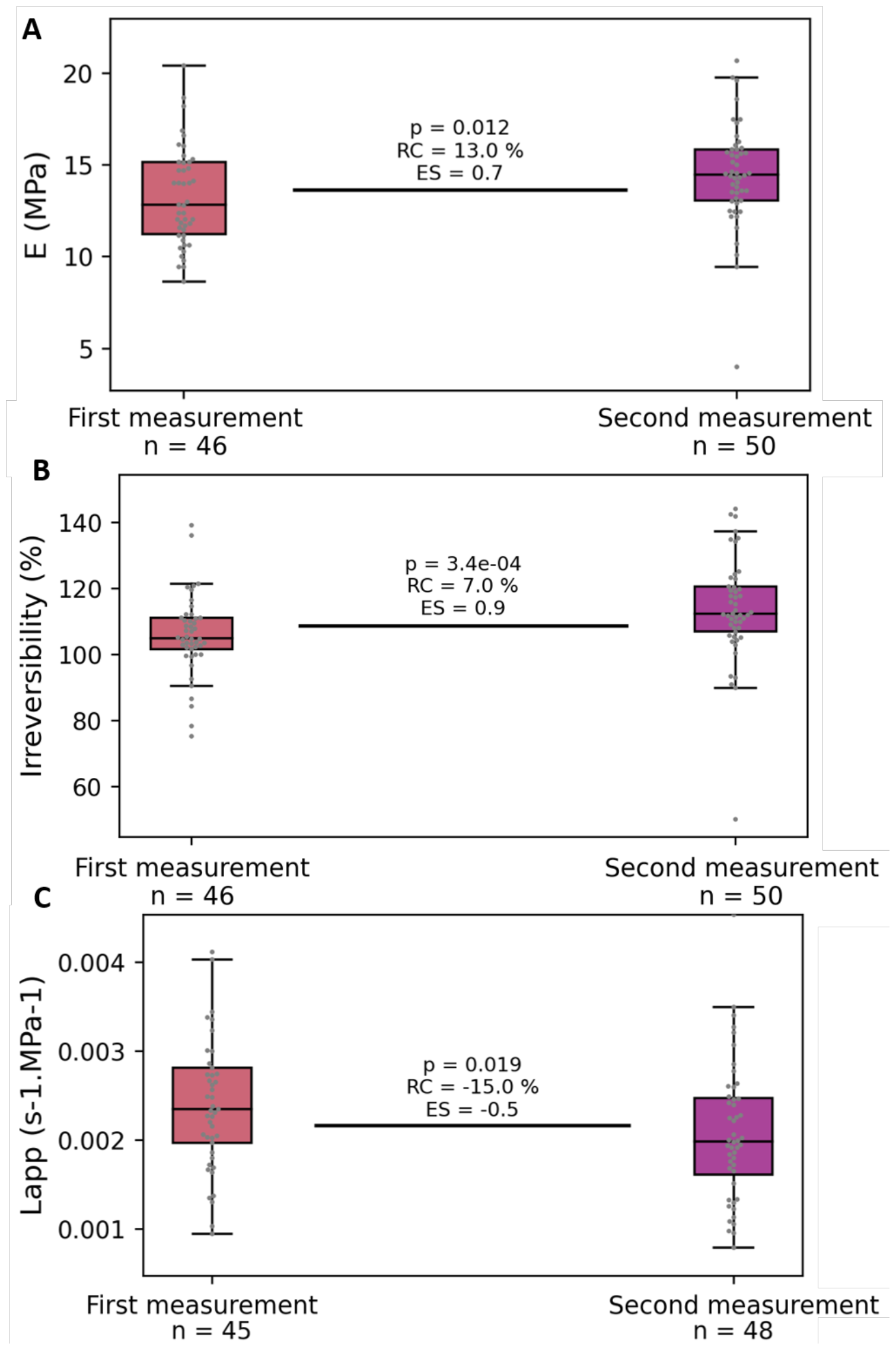
Consecutive osmotic steps. **A**. Comparison of successive bulk moduli measured at 30h and at 30h30min. **B**. Comparison of irreversibility levels measured in the same experiment as in (A). **C**. Comparison of apparent conductivity measured in the same experiment as in (A).

**Figure S3.**
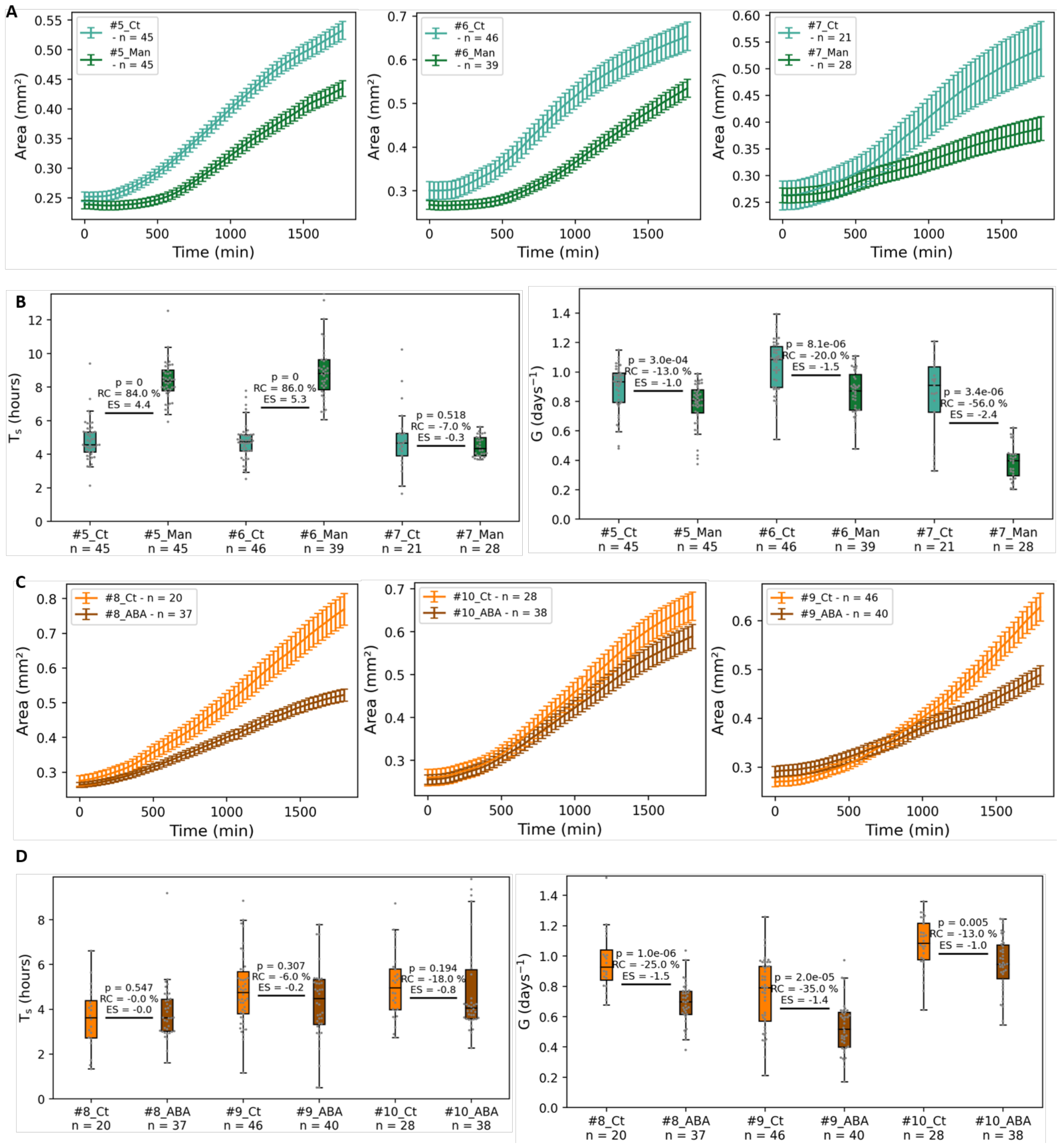
Growth in perturbed conditions – Detailed data. **A**. Growth curves per experiment for 100mM mannitol treatment (Man) and control (Ct) **B**. Quantification of the growth parameters corresponding to curves in (A) **C**. Growth curves per experiment abscisic acid treatment (ABA, 3.5µM) and control (Ct) **D**. Quantification of the growth parameters corresponding to curves in (C).

**Figure S4.**
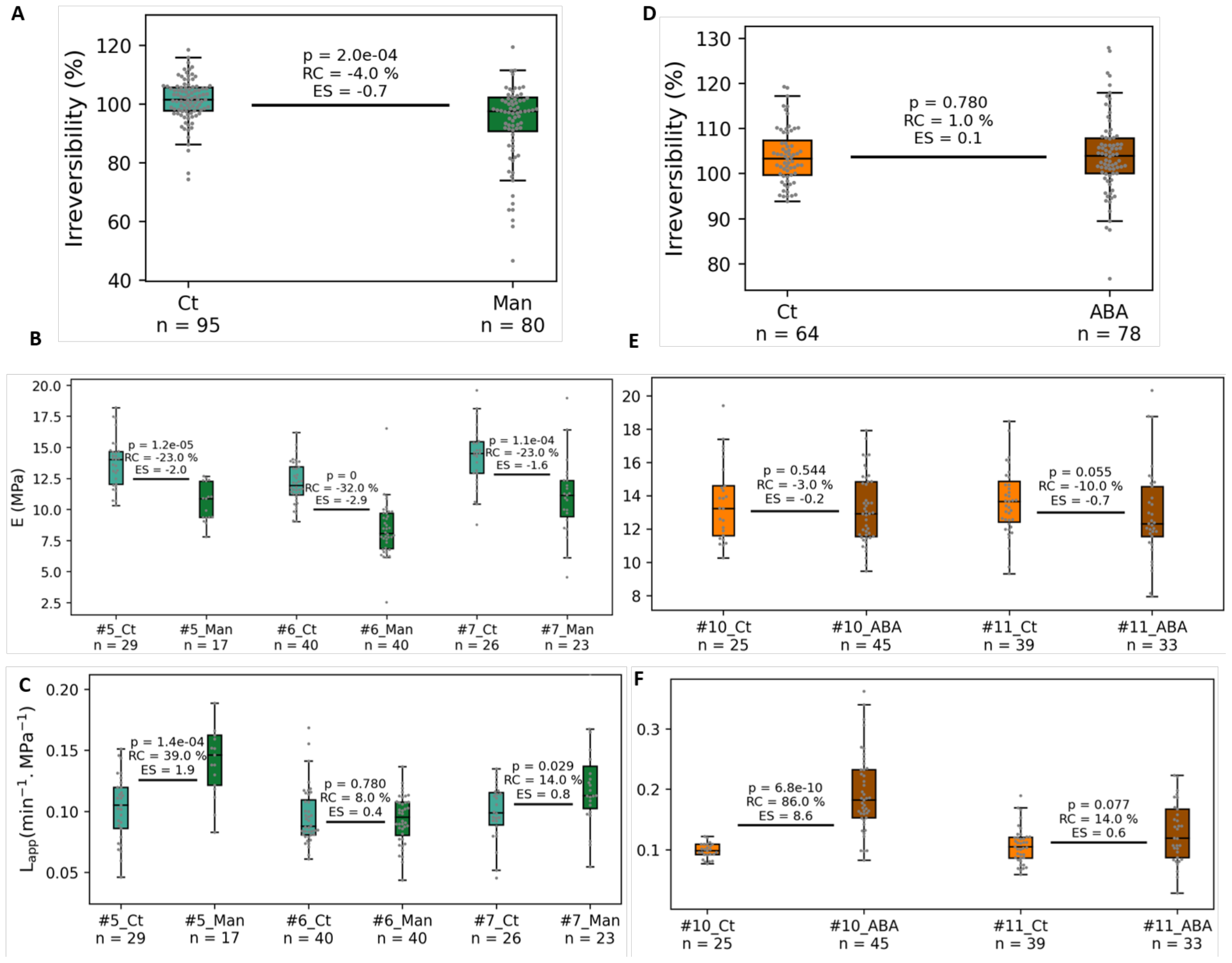
Hydromechanics in perturbed conditions – supplementary data. **A**. Irreversibility levels for the pooled experiments presented in Fig. 4-A: control and mannitol treated gemmae (Ct: control, Man: mannitol-treated). **B**. Detail of individual experiments pooled in Fig. 4-A. **C**. Detail of individual experiments pooled in Fig. 4-C. **D**. Irreversibility levels comparison for the pooled experiments presented in Fig. 4-C: control and ABA treated gemmae (Ct: control, ABA: abscisic acid-treated).. **E**. Detail of individual experiments pooled in Fig. 4-B. **F**. Detail of individual experiments pooled in Fig. 4-D.

**Figure S5.**
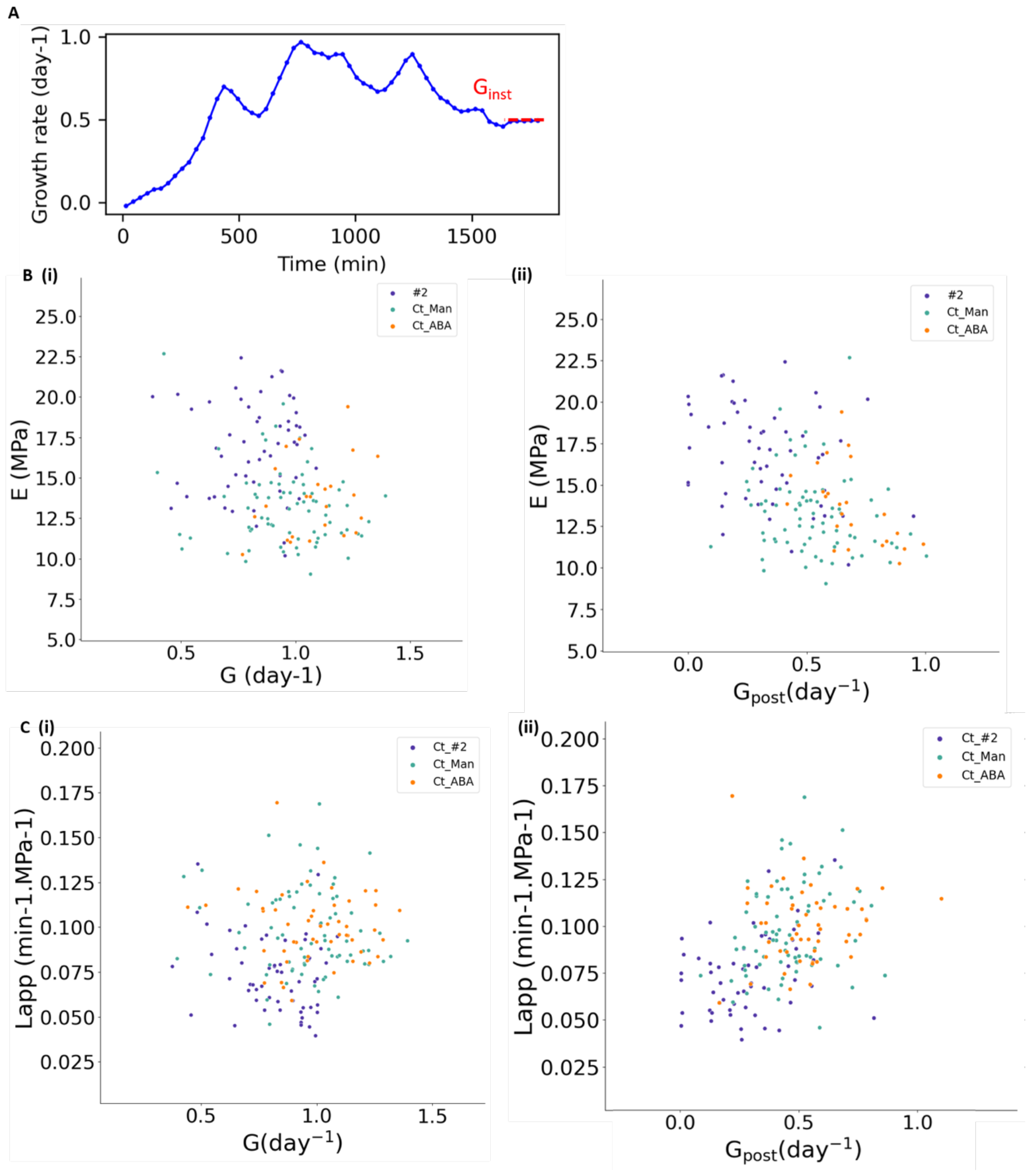
Link between hydromechanics and growth – Additional data. **A**. Computation of instantaneous growth rate G_inst_. **B**. Relation between the elastic modulus of gemmae and the average growth rate G (i, correlation coefficient CC = -0.197) as well as with the growth rate after the step G_post_ (ii, CC = -0.447). **C**. Relation between the apparent conductivity of gemmae and the average growth rate G (i, CC = 0.135) as well as with the growth rate after the step G_post_ (ii, CC = 0.431).

**Fig S6.**
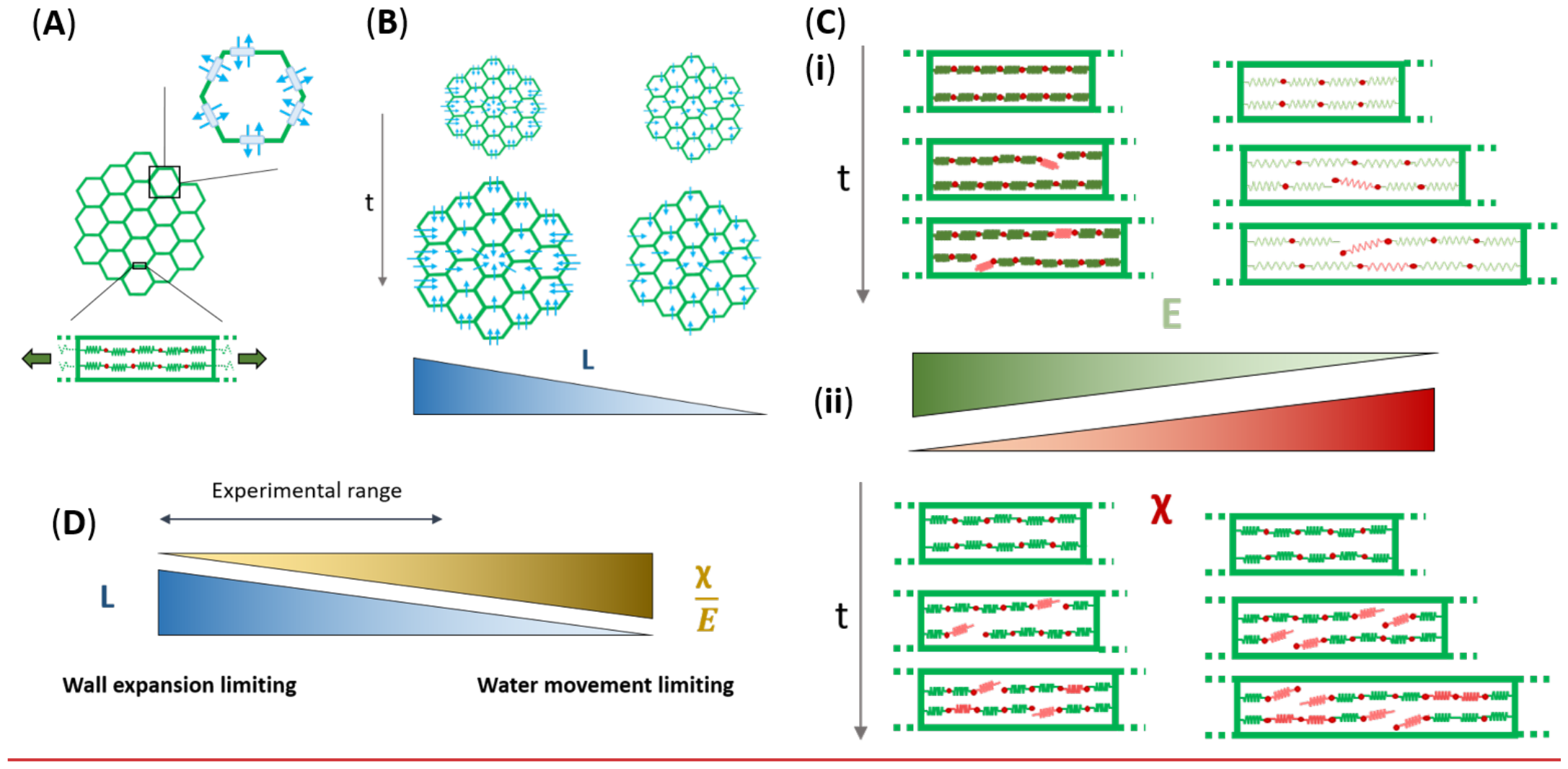
Model summary. **A**. Tissue growth may be limited by either water flow through the tissue (notably through pores: aquaporins and plasmodesmata) or by wall expansion (addition of new material, remodelling, wall elastic deformation). **B**. At high hydraulic conductivity, **L**, water can enter the tissue quickly, promoting fast growth (left). At low **L**, less water enters the tissue per unit time, slowing down growth (right). **C**. Expansion of the cell wall is regulated both by its ability to deform elastically (modulus E, part **i**) and by remodelling and addition of new material (chemical rate **χ**, part **ii**). (**i**) When **E** is high (left) the wall deforms less under tension when bonds are broken during remodelling. For small **E** (right), deformations are larger. (**ii**) For low **χ** (right) there is little remodelling and addition of new material per unit time, limiting growth. At higher **χ** (left) remodelling and addition of new wall material is faster and so is tissue growth. Newly added polymer segments are depicted in red. **D**. Overall, the balance between water flow and wall expansion allows for the fine tuning of the tissue growth rate. The range corresponding to experimental observations is indicated by a double arrow.

